# STAG: Species Tree Inference from All Genes

**DOI:** 10.1101/267914

**Authors:** D.M. Emms, S. Kelly

## Abstract

Species tree inference is fundamental to our understanding of the evolution of life on earth. However, species tree inference from molecular sequence data is complicated by gene duplication events that limit the availably of suitable data for phylogenetic reconstruction. Here we propose a novel method for species tree inference called STAG that is specifically designed to leverage data from multi-copy gene families. By application to 12 real species datasets sampled from across the eukaryotic domain we demonstrate that species trees inferred from multi-copy gene families are comparable in accuracy to species trees inferred from single-copy orthologues. We further show that the ability to utilise data from multi-copy gene families increases the amount of data available for species tree inference by an average of 8 fold. We reveal that on real species datasets STAG has higher accuracy than other leading methods for species tree inference; including concatenated alignments of protein sequences, ASTRAL & NJst. Finally we show that STAG is fast, memory efficient and scalable and thus suitable for analysis of large multispecies datasets.

## Introduction

The correct species tree is fundamental to understanding the diversity and history of life on earth. To infer species trees, researchers typically combine sequence data from sets of orthologous sequences (Jarvis, et al. 2014; Mao, et al. 2015; Ruhfel, et al. 2014). Often this is sets of protein coding genes, but can also include conserved non-transcribed elements as well as RNA genes and other nucleotide sequences. A common approach to integrating the information contained within multiple different genes is to join them together to form a concatenated multiple sequence alignment (CMSA). This is often preceded by checks to search for conflicts between the individual trees (James, et al. 2006; Perelman, et al. 2011; Ruhfel, et al. 2014) as well as tests to determine whether the data should be partitioned between genes (James, et al. 2006), data type (Jarvis, et al. 2014; Perelman, et al. 2011) or according to Akaike or Bayesian information criteria (Mao, et al. 2015; Meredith, et al. 2011). However, doubts have been raised about CMSA, with problems ranging from high bootstrap support for contentious branches (Salichos and Rokas 2013), to its statistical inconsistency under the multi-species coalescent model of incomplete lineage sorting (Roch and Steel 2015). Methods such as NJst (Liu and Yu 2011) and ASTRAL (Mirarab, et al. 2014) have been developed that bypass the concatenation problem and infer a species tree from a set of gene trees. However, both concatenation methods such as NJst and ASTRAL require data from groups of singlecopy orthologues genes, and the number of such genes in a group of species can be limited due to differing patterns of gene duplication and loss in the species sets being considered.

Species tree inference methods such as PHYLDOG (Boussau, et al. 2013) and Guenomu (Martins, et al. 2016) are not restricted to one-to-one orthologues and can therefore use more of the available whole-genome data. For example, PHYLDOG (Boussau, et al. 2013) jointly infers the species tree and the gene trees under a maximum likelihood model that combines a model for sequence evolution with a model for gene duplication and loss. However, the method is not suited to large datasets: to analyse 36 species and at most 100 genes per gene family, the method used 3000 processors of one of the top 500 supercomputers at the time (Boussau, et al. 2013). Thus, species tree inference methods that use single-copy genes are restricted in the amount of data they can access, and methods that can use multi-copy genes require computational resources that are beyond the reach of most research groups.

Here we present STAG (Species Tree inference from All Genes), a novel algorithm for inferring a species tree from sets of multi-copy gene trees. Through application to 12 real world datasets sampled throughout the eukaryotic domain we show that STAG is the most accurate method for species tree inference. Moreover, we demonstrate that consensus species trees generated using STAG have properties such as realistic support values and branch lengths that are suitable for downstream comparative analysis.

## Results

### Problem definition and benchmark datasets

Species tree inference methods are commonly tested on simulated sequence data (Liu and Yu 2011; Martins, et al. 2016; Mirarab, et al. 2014). However, the statistical similarity of these datasets to real biological datasets is not examined and the performance of the tested methods is not compared to expert curation of known species trees. To rectify this and provide comparative evaluation of species tree inference methods on real biological data, a collection of 12 diverse clades of species were sampled from throughout the eukaryotic domain (Table 1) (Emms and Kelly 2017). These datasets have varying numbers of species and rates of gene duplication (Emms and Kelly 2017), and have a broad range of estimated divergence times from c. 56 My for the Primates (dos Reis, et al. 2014) to c. 1,500 My for the Green Plants (Parfrey, et al. 2011). For each clade of species a published study was identified that provides a reference topology for the species tree derived from expert curation of molecular data (Table 2, Supplemental File S1). In all cases this reference topology is taken as true and is used as a benchmark for species tree inference method evaluation.

**Table 1:** Descriptive statistics of the 12 species datasets.

| Group | Species | Genes | Orthogroups | Orthogroups:<br>all species<br>present | Orthogroups:<br>all species<br>present &<br>single-copy | % Single-copy |
| --- | --- | --- | --- | --- | --- | --- |
| Birds | 47 | 708005 | 14454 | 4552 | 2616 | 57.5% |
| Flies (Diptera) | 7 | 120847 | 11688 | 3536 | 513 | 14.5% |
| Fish | 11 | 314941 | 16520 | 9585 | 518 | 5.4% |
| Fungi | 21 | 259240 | 9325 | 1583 | 338 | 21.4% |
| Hymenoptera | 5 | 80159 | 9157 | 7221 | 3947 | 54.7% |
| Kinetoplastids | 16 | 147588 | 9731 | 3317 | 1476 | 44.5% |
| Laurasiatheria | 14 | 274077 | 15804 | 6118 | 1386 | 22.7% |
| Metazoa | 21 | 407551 | 13017 | 1773 | 102 | 5.8% |
| Nematoda | 7 | 134567 | 8392 | 3187 | 517 | 16.2% |
| Primates + outgroup | 11 | 383525 | 19096 | 7142 | 286 | 4.0% |
| Rodents | 7 | 169136 | 15485 | 9296 | 2505 | 26.9% |
| Plants | 42 | 1366268 | 28356 | 1823 | 0 | 0.0% |

**Table 2:** Brief summary of method used for reference species tree inference. For full details see Supplemental File S1.

| Group | Species Tree Method |
| --- | --- |
| **Birds** | CMSA nucleotides, mixed, partitioned, third nucleotide of codons excluded |
| **Diptera** | CMSA nucleotides, genes, partitioned, third nucleotide of codon removed |
| **Fish** | CMSA, nucleotides, genes, partitioned by codon position |
| **Fungi** | CMSA, partitioned by gene (6), 3 AA, 3 RNA genes |
| **Hymenoptera** | CMSA, nucleotides, mixed, partitioned |
| **Kinetoplastids** | CMSA, amino acids |
| **Laurasiatheria** | CMSA, amino acids |
| **Metazoa** | CMSA, amino acids |
| **Nematoda** | MSA, SSU rRNA |
| **Primates** | CMSA, nucleotide, mixed, partitioned |
| **Rodents** | CMSA, amino acids |
| **Plants** | CMSA, nucleotides , partitioned |

**Table 3:**
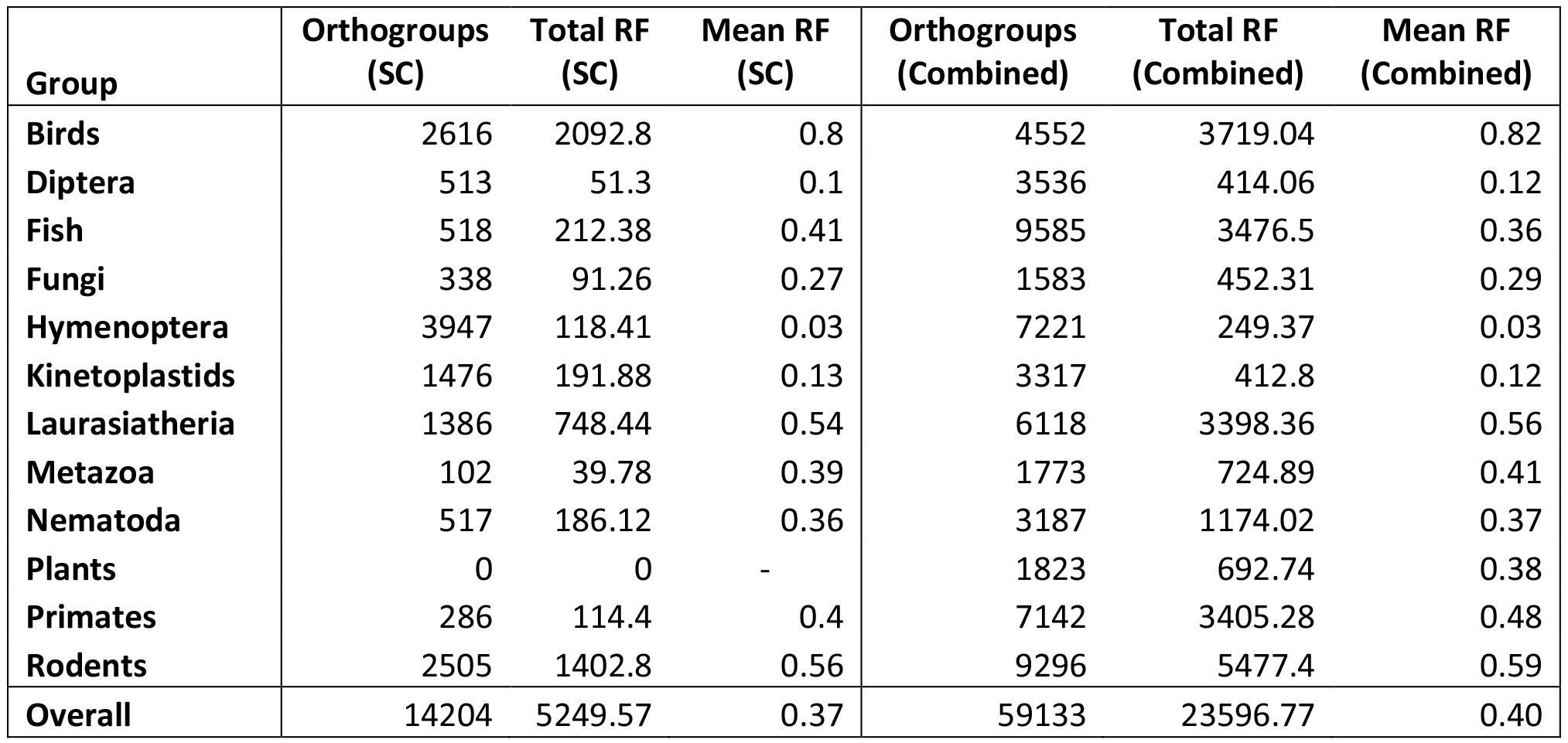
Comparison of the accuracy of trees inferred using single-copy and multi-copy genes.

| Group | Orthogroups (SC) | Total RF (SC) | Mean RF (SC) | Orthogroups (Combined) | Total RF (Combined) | Mean RF (Combined) |
| --- | --- | --- | --- | --- | --- | --- |
| Birds | 2616 | 2092.8 | 0.8 | 4552 | 3719.04 | 0.82 |
| Diptera | 513 | 51.3 | 0.1 | 3536 | 414.06 | 0.12 |
| Fish | 518 | 212.38 | 0.41 | 9585 | 3476.5 | 0.36 |
| Fungi | 338 | 91.26 | 0.27 | 1583 | 452.31 | 0.29 |
| Hymenoptera | 3947 | 118.41 | 0.03 | 7221 | 249.37 | 0.03 |
| Kinetoplastids | 1476 | 191.88 | 0.13 | 3317 | 412.8 | 0.12 |
| Laurasiatheria | 1386 | 748.44 | 0.54 | 6118 | 3398.36 | 0.56 |
| Metazoa | 102 | 39.78 | 0.39 | 1773 | 724.89 | 0.41 |
| Nematoda | 517 | 186.12 | 0.36 | 3187 | 1174.02 | 0.37 |
| Plants | 0 | 0 | - | 1823 | 692.74 | 0.38 |
| Primates | 286 | 114.4 | 0.4 | 7142 | 3405.28 | 0.48 |
| Rodents | 2505 | 1402.8 | 0.56 | 9296 | 5477.4 | 0.59 |
| Overall | 14204 | 5249.57 | 0.37 | 59133 | 23596.77 | 0.40 |

### Sets of one-to-one orthologues are rare, and decrease as a function of cumulative evolutionary distance between sampled species

To provide a common input dataset on which to test multiple species tree inference methods orthogroups were inferred for each of these 12 datasets using OrthoFinder (Emms and Kelly 2015). Several of the methods under consideration here require datasets of one-to-one orthologues that are present in all (or most) of the species being investigated. Thus the number of genes per species in each orthogroup was analysed. For each species set, there are large numbers of orthogroups in which every species is present. However, few of these orthogroups contain just one orthologue from each species (single-copy orthogroups, Figure 1). On average there are 8.4 times more orthogroups with all species present than single-copy orthogroups with all species present (Table 1). In the plant species dataset large numbers of gene and genome duplication events (Lee, et al. 2013) mean that there are no single-copy orthogroups present in all species (Table 1). In contrast, at the other end of the spectrum the bird species have small genomes (Jarvis, et al. 2014) and low rates of gene duplication (Emms and Kelly 2017) resulting in large numbers of single-copy orthogroups (Table 1). Thus in real world datasets, while orthogroups that contain all species are common, orthogroups that contain just one orthologue from each species comparatively rare.

**Figure 1.**
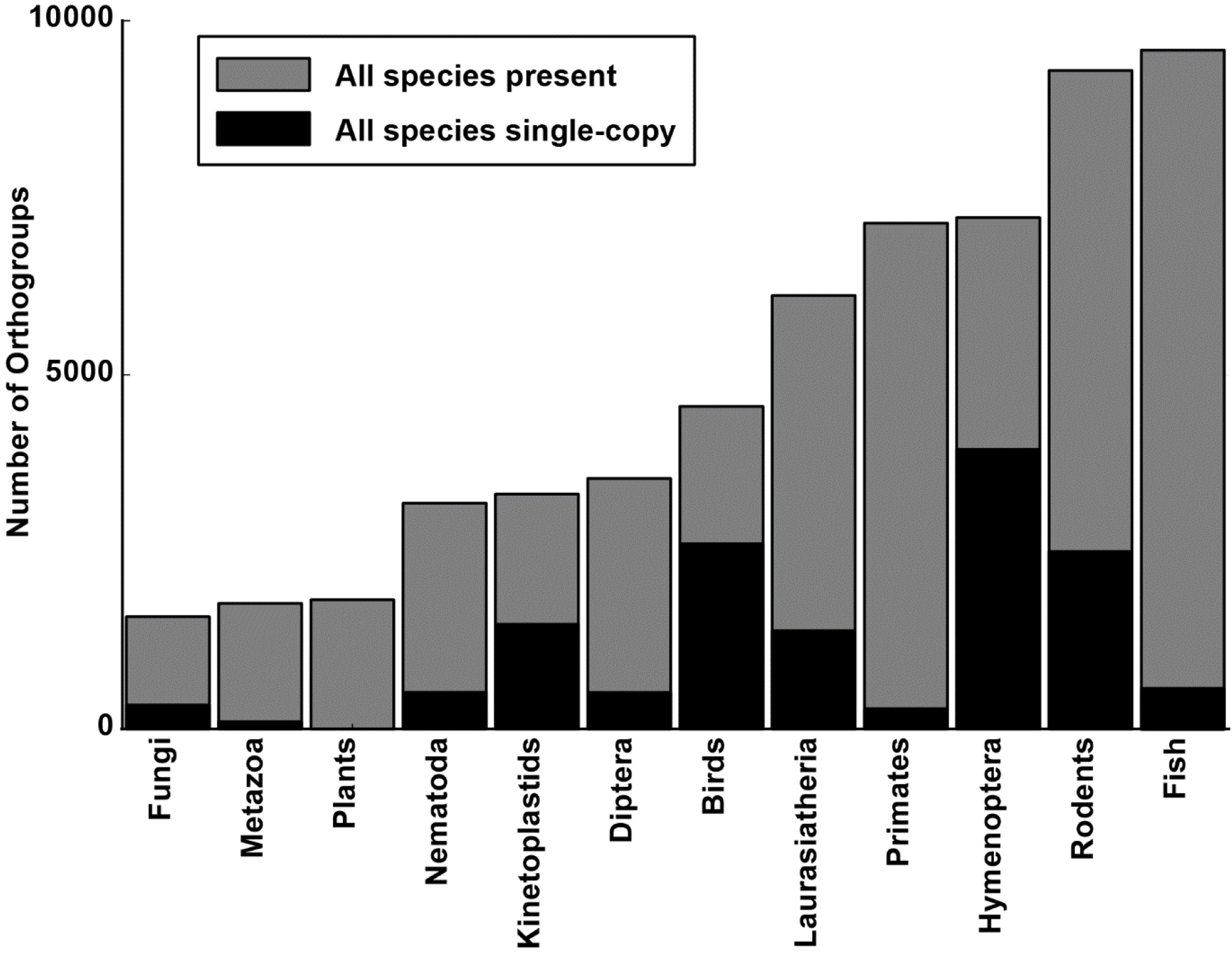
The number of orthogroups with all species present (grey bars) and with all species present and single-copy (black bars) in the 12 biological datasets.

Across the 12 biological datasets the number of single-copy orthogroups was negatively correlated with the number of species under consideration, but not significantly so (Figure 2A, r^2^=0.22, p = 0.12). Taking divergence time into consideration too, the number of single-copy orthogroups was significantly negatively correlated with the species tree length (Figure 2B, r^2^=0.62, p = 0.002). Thus, the number of single copy orthogroups available for analysis decreases due to the combined effect of increasing number of species and the divergence time of those species.

**Figure 2.**
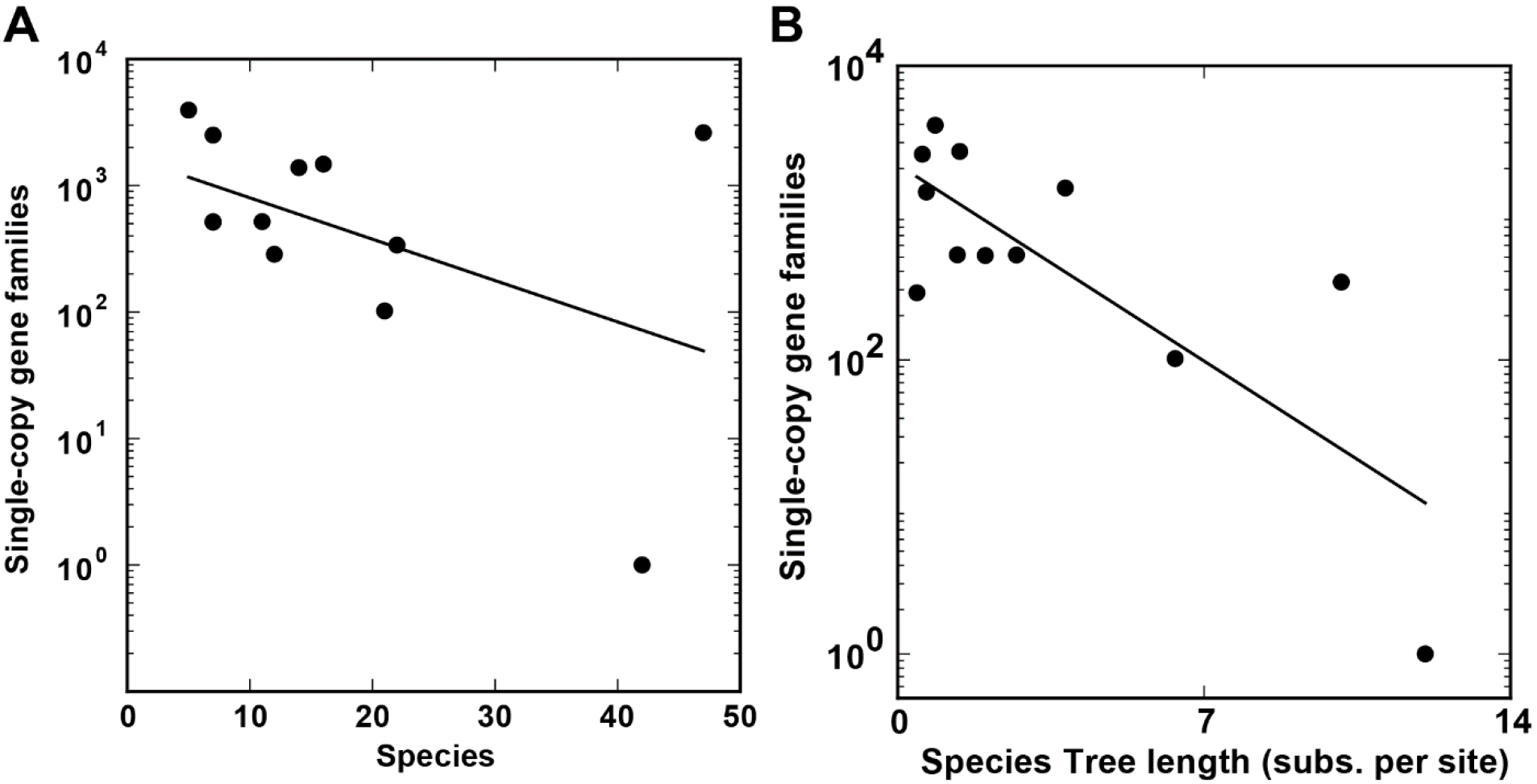
The number of orthogroups with all species present and single-copy in the 12 biological datasets versus A) the number of species in that dataset B) The species tree length.

### Species trees from multi-copy gene families are of comparable accuracy to those from single-copy gene families

Given the limited availability of single copy orthogroups in real biological datasets described above, the ability to use multi-copy orthogroups would help mitigate the problem of low data availability. However, existing methods that can use multi-copy genes require computational resources that are beyond the reach of most research groups (Boussau, et al. 2013). To address this problem, a novel method called STAG (Species Tree from All Genes) was developed. Full details of the algorithm are provided in the Methods. In brief, STAG analyses each multi-copy gene tree in turn. For a given tree, the distances between each pair of sequences on the gene tree are analysed. For a given species pair, the gene-pair with the shortest distance on the tree is selected as these genes will most likely be orthologues. In the unlikely event where these gene pairs are not orthologues, then the selected gene pair is more closely related than any pair of orthologues for the same species-pair in the same gene tree. A species tree is then inferred from this gene tree by evaluating the set of minimum pairwise distance estimates between all species using the minimum evolution principle (Lefort, et al. 2015). In this way each multi-copy gene tree is able to provide an estimate of the underlying species tree.

To test the accuracy of these species trees inferred from multi-copy genes, the complete set of species trees inferred from multi-copy gene trees from the 12 species sets were subject to topological analysis. Here, each multi-copy gene orthogroup was subject to multiple sequence alignment using MAFFT L-INS-i (Katoh and Standley 2013) and phylogenetic inference using IQ-TREE (Nguyen, et al. 2015). Species trees were inferred from each multi-copy orthogroup gene tree using STAG, and the Robinson-Foulds distance (Robinson and Foulds 1981) between the STAG tree and the published species tree was evaluated. To place these results in context, the complete set of gene trees inferred by IQ-TREE from single copy orthogroups for the same species dataset was also subject to the same Robinson-Foulds distance analysis. Single copy gene trees produced in this manner are conventionally used for species tree inference as they each contain an estimate of the underlying species tree. Comparison of the two approaches revealed that species trees inferred from multi-copy gene orthogroups using STAG were of comparable accuracy to species trees inferred from single copy gene orthogroups using conventional methods (Figure 3, Supplemental File S1 Table S1).

**Figure 3.**
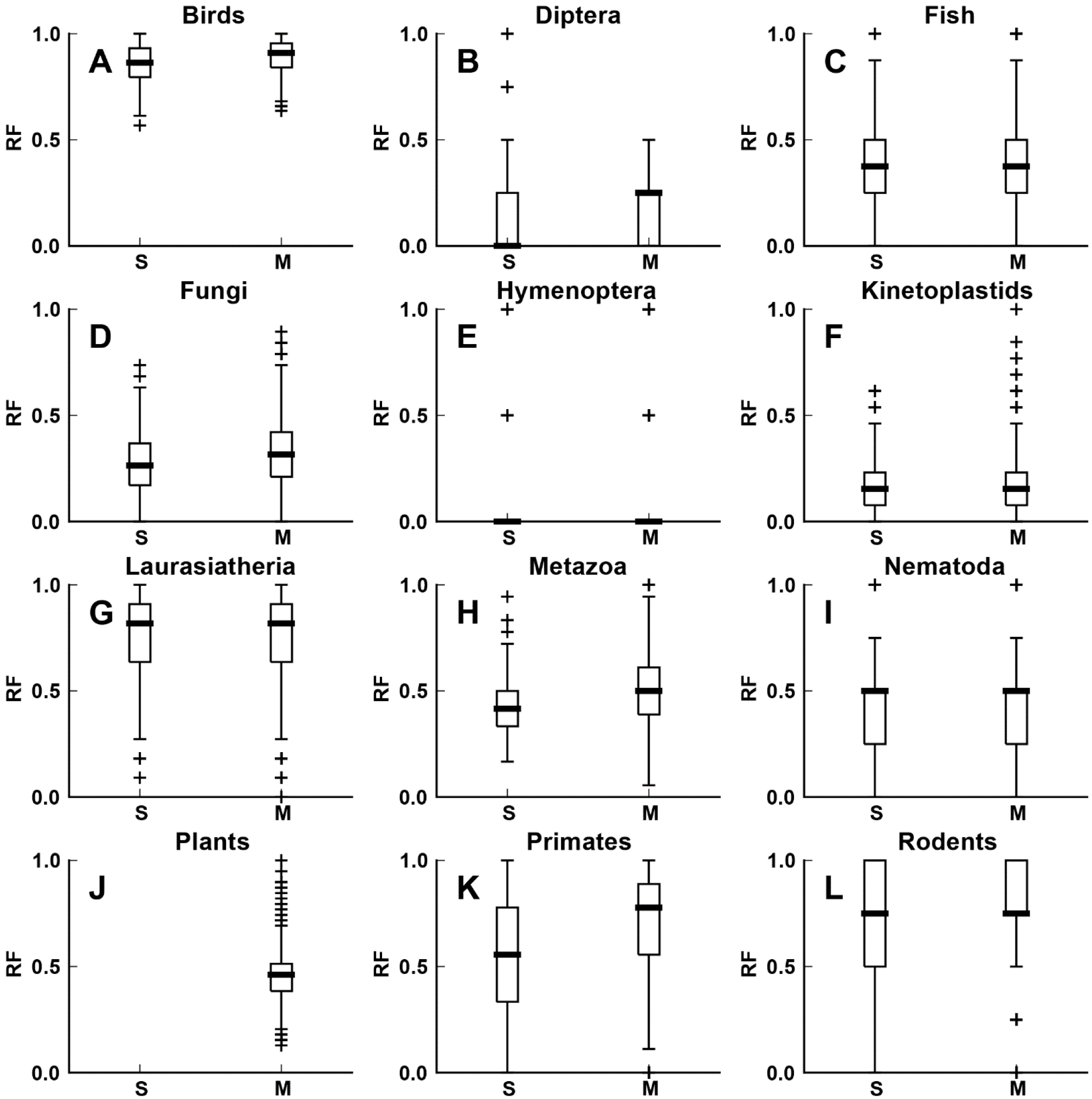
The distribution of Robinson-Foulds distances between the literature-derived species tree and i) the IQ-TREE trees from orthogroups with all species present and single-copy (S) ii) the individual per-orthogroup species trees inferred by STAG from orthogroups with all species present but not singlecopy in all species (i.e. multi-copy genes M). Results are for the 12 biological datasets (A-L), for the plants dataset there were no orthogroups identified with all species present and single-copy and thus there is no data for S.

Across the 12 biological datasets, inclusion of STAG trees extracted from multi copy gene orthogroups increased the mean RF distance of the set of trees available for species tree inference by ~3% (Table 2). However, the average amount of data available for species tree inference increased on average by 800% (Table 2). Thus, for a small increase in the amount of error per tree, there is a substantial increase in the data available for tree inference.

### Consensus species trees inferred using STAG are more accurate than other methods on real biological datasets

Given STAG provides a substantial increase in data availability with only a small penalty to mean dataset accuracy it was determined how this would affect the overall accuracy of species tree inference when these trees are integrated. To investigate this, a consensus species tree for each of the 12 species datasets was inferred by taking the greedy consensus (Felsenstein 2005; Swofford and Sullivan 2009) of each estimate of the STAG trees inferred from the sets of orthogroups with all species present. Thus, consensus STAG trees contain support values at internal bipartitions that quantify the proportion of input trees in which that bipartition occurs. The number of input trees used for species inference for each group is given in Table 1; the number of orthogroups with all species present.

To place these results in context, species trees for the same species datasets were also inferred with a number of leading methods for species tree inference (Supplemental File S1 Table S2). This set comprised ASTRAL (Mirarab and Warnow 2015), Concatenated MSA (CMSA), NJst (Liu and Yu 2011) and Guenomu (Martins, et al. 2016). Each method was run using best practice approaches for these methods (see Methods). The species tree produced by STAG agreed with the published species tree more frequently than any of the other methods (Figure 4, Table 4). Moreover, the median and mean Robinson-Foulds distance between the STAG consensus species tree and the published species tree was lower than for any other method (Figure 4, Table 4). An example tree where all of the tested methods disagree is provided in Figure 5. Here, partitions that are incorrect in the STAG consensus species tree receive low support values in contrast to the other tested methods (Figure 5).

**Figure 4.**
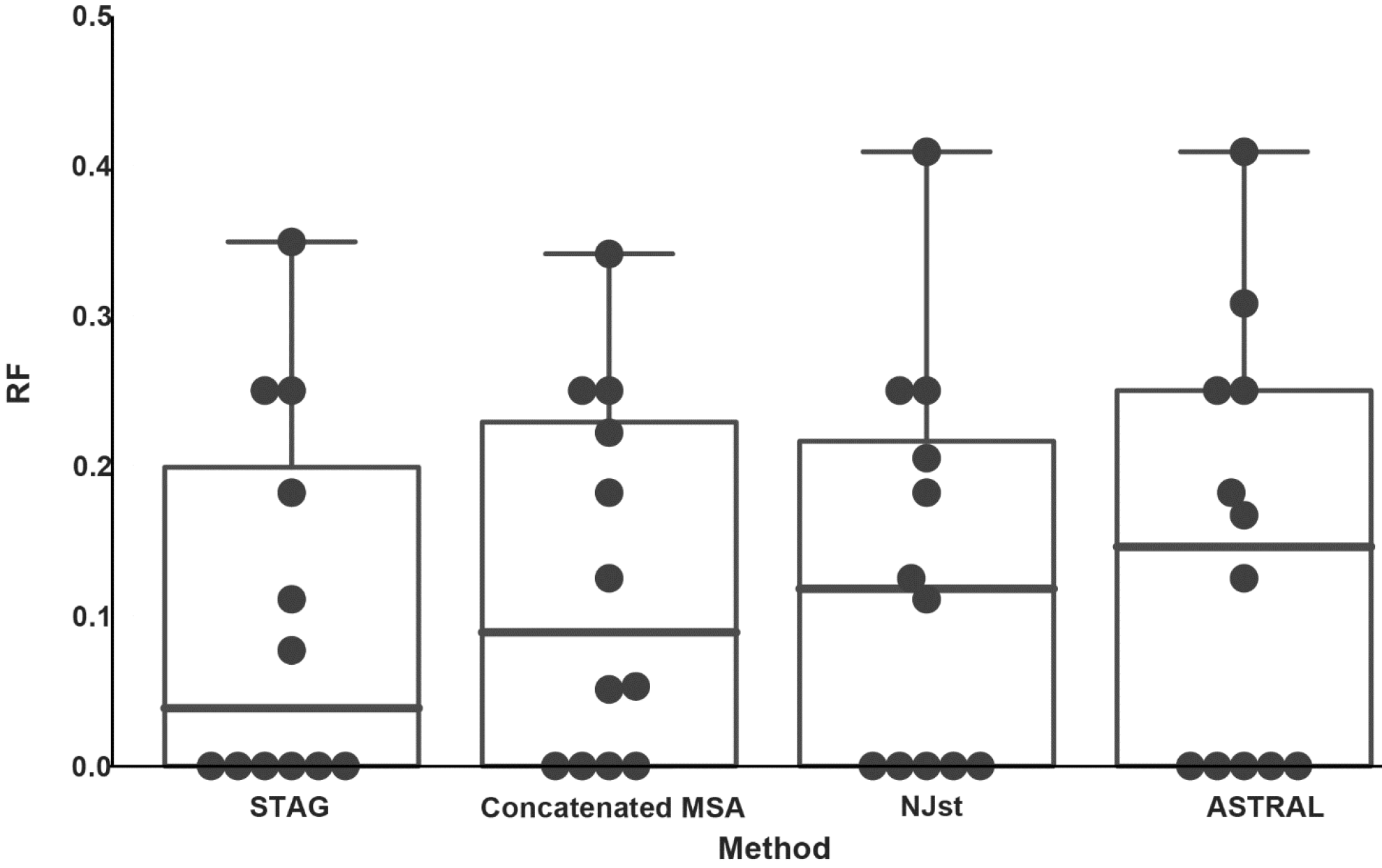
Box plots for the Robinson-Foulds (RF) distance between the literature-derived species tree and the species trees inferred by each of the tested methods across the 12 biological datasets. The dots give the individual RF distances for each of the 12 datasets.

**Figure 5.**
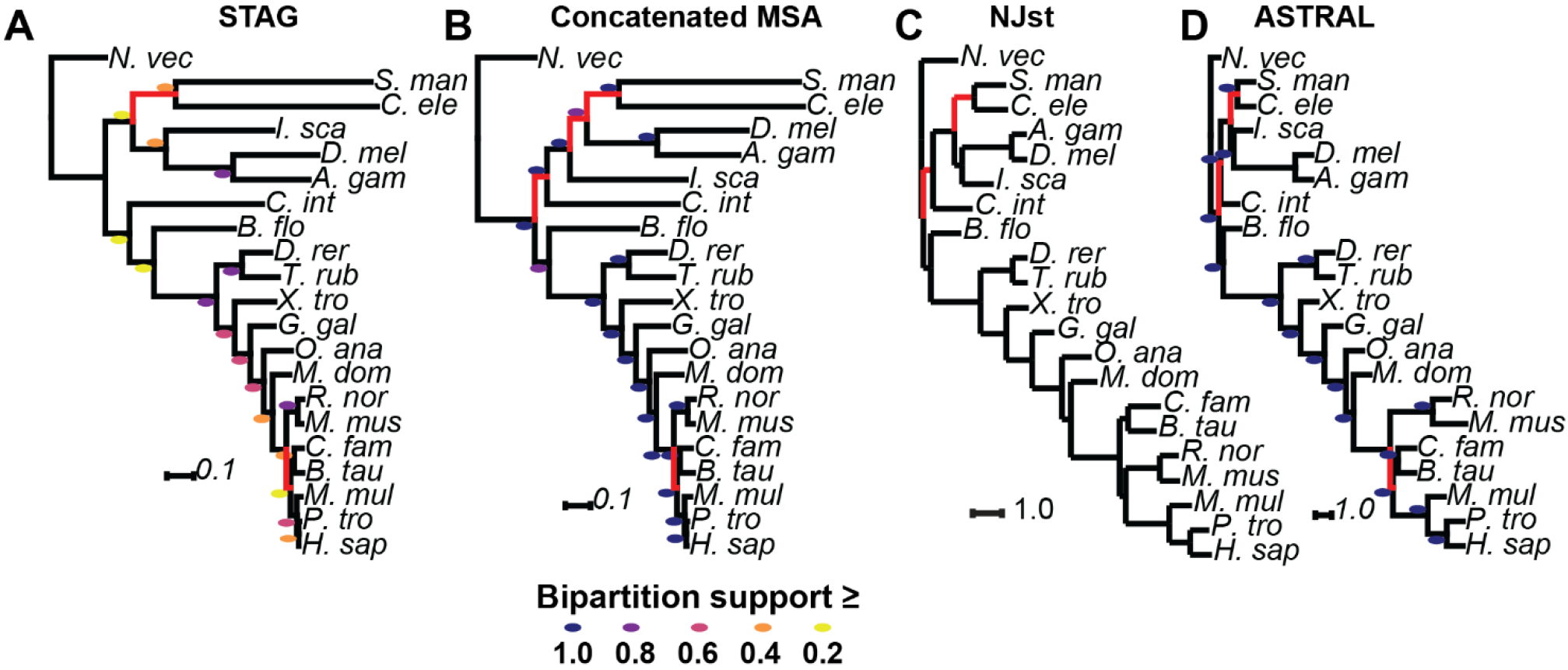
The species trees inferred for the metazoa dataset by A) STAG B) CMSA C) NJst D) ASTRAL. Branch lengths are in A-B) substitutions per site or C-D) coalescent units. Bipartition support values are colour-coded next to each branch for those methods returning support values. A scale for the colour coding is provided in the figure.

**Table 4:** Evaluation of species tree inference methods on 12 benchmark datasets.

| Clade | STAG | CMSA | NJst | ASTRAL | Guenomu |
| --- | --- | --- | --- | --- | --- |
| Birds | 0.349 | 0.341 | 0.409 | 0.409 | Timeout |
| Diptera | 0 | 0 | 0 | 0 | 0 |
| Fish | 0.25 | 0.125 | 0.125 | 0.125 | 0.125 |
| Fungi | 0 | 0.053 | 0 | 0 | 0.474 |
| Hymenoptera | 0 | 0 | 0 | 0 | 0 |
| Kinetoplastids | 0 | 0 | 0 | 0 | 0 |
| Laurasiatheria | 0.182 | 0.182 | 0.182 | 0.182 | 0.182 |
| Metazoa | 0.111 | 0.222 | 0.111 | 0.167 | No result |
| Nematoda | 0.25 | 0.25 | 0.25 | 0.25 | 0.25 |
| Plants | 0.077 | 0.051 | 0.205 | 0.308 | Timeout |
| Primates | 0 | 0 | 0 | 0 | No result |
| Rodents | 0 | 0.25 | 0.25 | 0.25 | 0 |
| Mean | **0.102** | **0.123** | **0.128** | **0.141** | - |

### Branch lengths obtained from STAG consensus trees are comparable to those obtained concatenated multiple sequence alignments

Given that STAG performed well on the tree topological tests described above it was investigated whether the branch lengths of the STAG consensus trees accurately represented molecular phylogenetic distances between species. To provide an unbiased comparison, only the species trees where all methods were correct and thus had 100% agreement on topology were analysed (Figure 6). The branch lengths obtained from STAG consensus trees correlate well with those produced by other methods (mean r^2^ = 0.72), and are essentially identical to those produced using concatenated multiple sequence alignments (r^2^ = 0.99, p = 10^−77^, Figure 6 & Supplemental File S1 Table S3). Thus branch lengths provided by STAG consensus trees are suitable for use in downstream analyses such as ancestral state reconstruction or time calibration.

**Figure 6.**
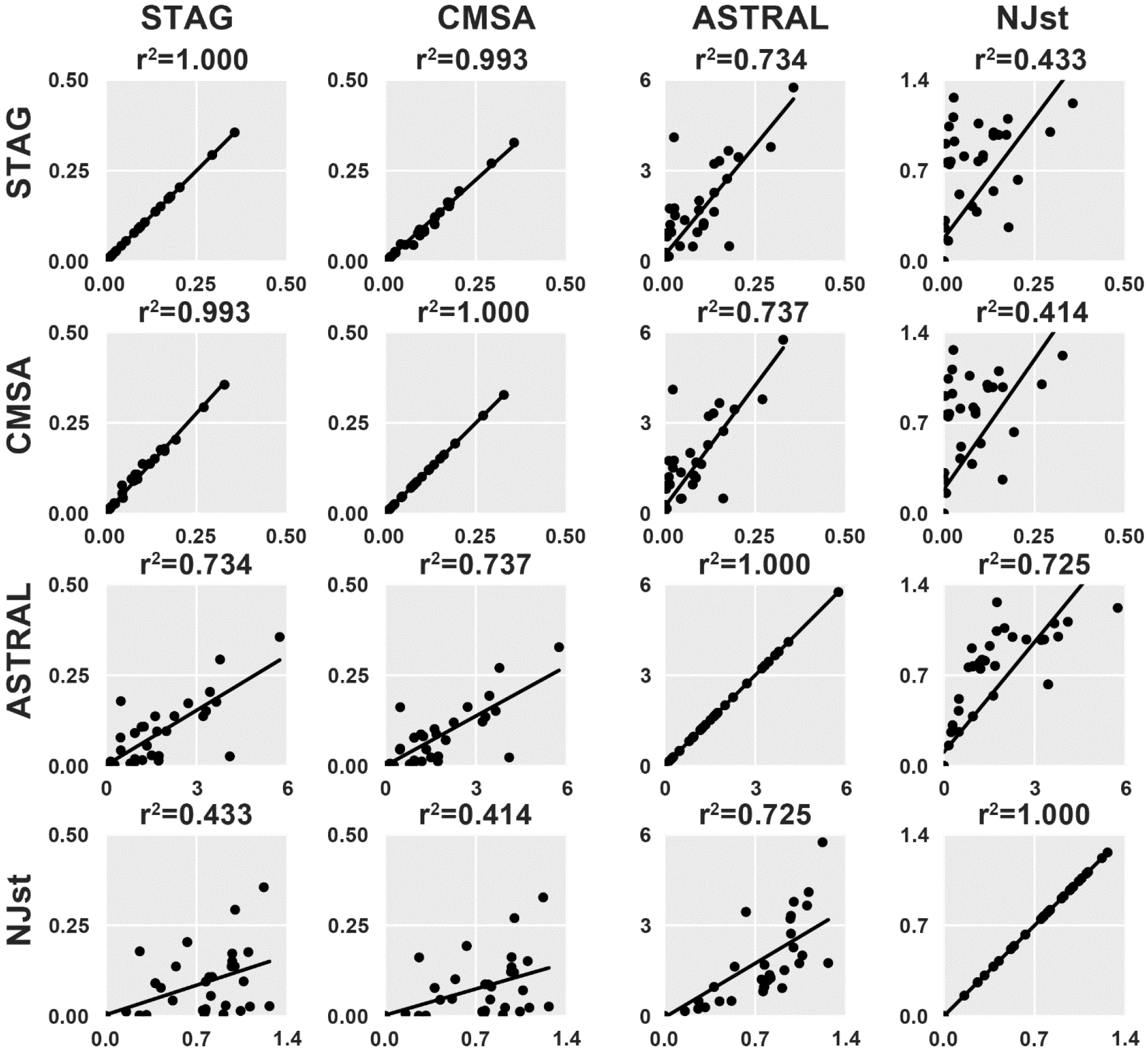
Correlation between the branch lengths returned by STAG, CMSA, ASTRAL & NJst for the datasets for which all four methods returned the same (correct) species tree topology. Line of best fit is for the linear least-squares regression, with the r^2^ value given above each plot.

### STAG is fast and efficient

To demonstrate the performance characteristics of STAG the time and RAM usage across the 12 species datasets was analysed. The maximum time and RAM usage for STAG to infer a consensus species tree on a single core of a conventional desktop computer was 95.1 seconds and 0.12 GB respectively (Table 5). Across the 12 datasets the run time was linearly dependent on the number of species and number of trees being analysed (Supplemental File S1 Figure S1). Similar the RAM requirements were linearly dependent on the number of species (Supplemental File S1 Figure S1).

**Table 5:** Performance characteristics of STAG.

|  | Species | Trees | Time<br>(s) | RAM<br>(GB) |
| --- | --- | --- | --- | --- |
| Birds | 47 | 4553 | 95.1 | 0.12 |
| Diptera | 7 | 3536 | 17.2 | 0.05 |
| Fish | 11 | 9585 | 61.3 | 0.05 |
| Fungi | 21 | 1583 | 18.2 | 0.05 |
| Hymenoptera | 5 | 7220 | 29.1 | 0.04 |
| Kinetoplastids | 16 | 3317 | 24 | 0.05 |
| Laurasiatheria | 14 | 6118 | 44 | 0.05 |
| Metazoa | 21 | 1773 | 47.9 | 0.06 |
| Nematoda | 7 | 3187 | 16.6 | 0.04 |
| Plants | 42 | 7142 | 82.6 | 0.07 |
| Primates | 11 | 9296 | 54.2 | 0.05 |
| Rodents | 7 | 1823 | 47.9 | 0.05 |

## Discussion

With fully sequenced genomes available for many species, there is abundant sequence data from which to infer species trees. However, the majority of species tree inference methods are restricted to use one-to-one orthologous sequences that are present in all species in the analysis. Such groups of sequences are available only if gene duplication or loss has not occurred during the divergence of that gene family. In this work it is shown that such one-to-one orthogroups are comparatively rare in real biological datasets, and become rarer as species tree length increases (a product of increased divergence time and increased species sampling). We presented a novel method called STAG (Species Tree inference from All Genes). The method constructs a consensus species tree from trees from all orthogrous in which all species are present, irrespective of the gene copy number per-species in the orthogroup. On real species datasets STAG out-performed species trees inferred by comparable methods such as ASTRAL, NJst, and Guenomu as well as maximum likelihood trees inferred from concatenated multiple sequence alignments (CMSA).

The testing was performed using 12 real biological datasets sampled from throughout the eukaryotic domain. At the time of writing, this is the largest collection of biological datasets for testing species tree inference that has been assembled. The use of real biological datasets in species tree inference evaluation eliminates modelling assumptions that are required in order to generate simulated test datasets. Moreover, widely sampled real biological datasets reflect the true disparity in real species datasets and thus accurately represent the kinds of datasets to which the method will be applied. The disadvantage of biological data is that the ground truth is not known. Thus, for each of these 12 clades of species, a published study that inferred a species tree from expert curation was accepted as true in order to facilitate comparison and benchmarking of the methods tested here. The tests should not bias the results in favour of any of the tested species tree inference methods as they were not used to generate the references trees. Moreover, these tests should not bias towards STAG as multi-copy gene trees were not used in any of these studies. Furthermore, the 12 datasets presented here represent a significant increase over previous analyses that only used 1 (Liu and Yu 2011; Martins, et al. 2016) or at most 3 (Mirarab, et al. 2014) biological datasets.

STAG has been implemented as a freely available, standalone program. It has been designed to be easy to use. It requires as input a directory of gene trees (which do not have to be rooted) and a file describing how gene names map to species. It doesn’t require any pre-processing of the gene trees, for example to exclude trees with duplications or trees with too few taxa. It has been designed to integrate with OrthoFinder (Emms and Kelly 2015), which is a method for inferring the set of orthogroups for all genes in a set of species. It can be launched with a single command directly from a set of OrthoFinder results. The method is also fast, taking only 95.1 seconds on a single core on the largest dataset. The peak RAM usage was 0.12 GB. This analysis was for a set of 47 species and involved the inference of 4553 individual estimates of the species tree, which were combined to give the final STAG species tree. The gene trees that were used were all automatically generated and involved no expert curation.

Although STAG is fully automated and intended for use with large datasets, it is equally well-suited to the inference of the species tree from a carefully curated set of gene trees, as is common for studies that aim to resolve challenging clades of the tree of life. Thus careful filtering and processing assemblages of multi-gene family alignments and trees prior to running STAG will likely aid in increasing the accuracy of species tree inference. In this way, STAG will aid expert curation and analysis of phylogenetic datasets enabling substantial increases in data availability.

The units for the branch lengths in the STAG species tree are the same as the units in the input gene trees. In most cases, these will be the number of substitutions per site. The tree branch lengths in the STAG consensus tree are the average branch lengths across all the individual trees inferred from each gene family. Thus, the branch lengths represent the average number of substitutions per sites across a large range of gene families. This is an important feature of STAG trees, as these branch lengths can be used directly in downstream analyses such as ancestral state reconstruction and time calibration. Equivalent utility is not present in species trees generated using ASTRAL, NJst, or Guenomu. Furthermore, the support values for each bipartition in a consensus STAG tree are the proportion of times that the bipartition is seen in each of the individual species tree estimates. They are therefore lower, in general, than those given by the concatenation method, which can quickly reach 100% support even for bipartitions believed to be incorrect (Salichos and Rokas 2013) see also Figure 5.

## Materials and methods

### Algorithm overview

STAG obtains an estimate for divergence between each species pair from orthologous gene pairs in a given gene tree. Within a gene tree these estimates do not need to come from the same ortholog, but rather the closest estimate for each species pair is taken to mitigate against problems such as hidden paralogy. The input to STAG is a set of unrooted gene trees (Figure 7). For each gene tree containing all species, STAG identifies the closest pair of genes from those species as those that are separated by the shortest branch length. These shortest distances are used to construct an interspecies distance matrix, and a tree is inferred from this distance matrix using FastME (Lefort, et al. 2015). Thus, for each gene tree containing all species, an estimate is made of the underlying species tree. These individual estimates are combined using a standard greedy consensus method (Felsenstein 2005; Swofford and Sullivan 2009). The support for each bipartition in the STAG consensus species tree is equal to the proportion of individual estimates of the species tree that contain this bipartition. The branch lengths in the STAG consensus species tree are the average branch lengths for each bipartition in the individual estimates of the species tree. It is possible that in some individual estimates of the species tree, the gene pairs used to estimate inter-species distance are paralogues descended from a gene duplication event followed by the differential loss of the orthologue for each duplicate. This is known at hidden paralogy (Martin and Burg 2002) and is a problem that affects all of the other methods tested here and is thus not specific to STAG. The assumption that one-to-one genes in a tree are orthologues is common to all methods that infer trees from presumed orthologues.

**Figure 7.**
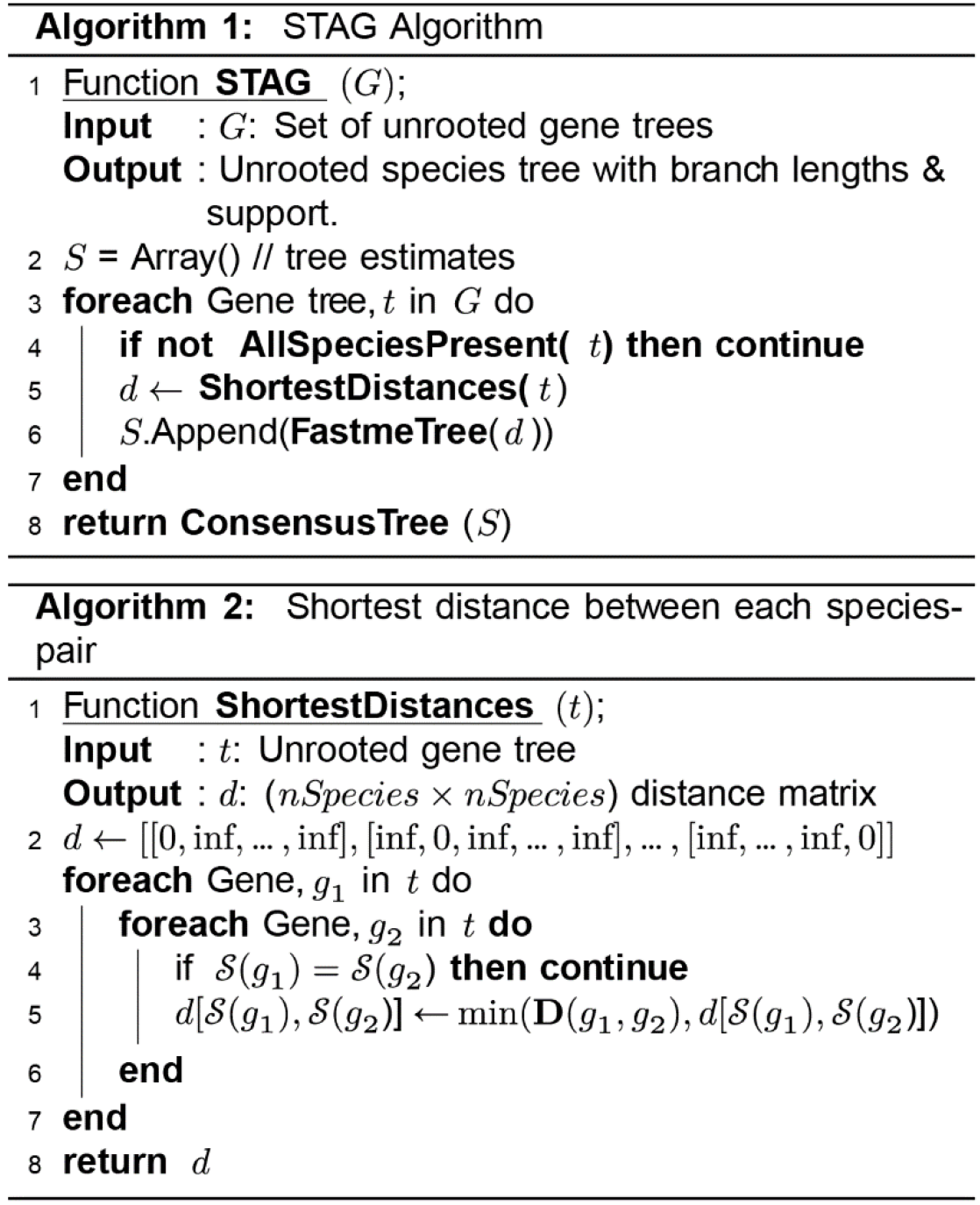
Pseudo-code for the STAG algorithm. The algorithm makes use of a number standard routines: ‘AllSpeciesPresent’ returns True if all species are present in a gene tree and False ortherwise; ‘FastmeTree’ calls the FastME program to calculate a tree from a distance matrix; ‘ConsensusTree’ calculates the greed consensus tree from a set of trees.

### Datasets for evaluation and benchmarking

We used the 12 biological datasets and literature-derived species tree topologies from a previous study (Emms and Kelly 2017). This consisted of a diverse set of species sampled from throughout the eukaryotic domain. This included every named group of eukaryotes on Ensembl Genomes containing >4 genera as well as sets of 47 Birds, 42 Green Plants and 16 Kinetoplastids. For each dataset, the species tree topology was taken the best available from a published study (Table 2, Supplemental File S1).

Orthogroups were inferred for each species set using OrthoFinder (Emms and Kelly 2015). Multiple sequence alignments (MSA) were inferred for each orthogroup using MAFFT L-INS-i (Katoh and Standley 2013) and gene trees were inferred from these MSAs using IQTree (Nguyen, et al. 2015). Appropriate subsets of this data were used to evaluate all methods presented in this study according to best practices described the following section.

### Implementation of comparative methods

There is no consensus best practice for construction of concatenated multiple sequence alignments for species tree inference (Supplemental File S1). To infer the CMSA species tree, each alignment was trimmed to include only those columns present in 50% or more of the species. The species tree was inferred from the concatenated alignment using IQ-TREE, as was done for the individual gene trees. We used ASTRAL version 5.5.9 with default parameters. We used the implementation of NJst available at https://github.com/adamallo/NJstM (last updated Dec 2016) using the original method. We used Guenomu version 201308. For Guenomu we used the provided control file, but at the authors suggestion reduced ‘param_reconciliation_prior’ to 10-9 to attempt to resolve the lack of convergence for some datasets. The Metazoa and Primates datasets returned a flat posterior distribution for the species tree and were recorded as ‘No result’ (equal probabilities for each of the sampled species tree topologies) To attempt to run the Birds and Plants datasets, we excluded the largest 200 orthogroups and reduced the number of number of sample generations and number of samples by a factor of 10 (to ‘param_n_generations = 5000 10000’ and ‘param_n_samples = 100’). We ran these datasets on the University of Oxford HPC ARCUS-B in parallel with 16 cores but they timed out at 120 hours without completing the initial 5000 burn-in generations (for comparison the longest runtime for STAG was 95.1s on a single core on a desktop machine). Thus the bird and plant datasets were recorded as ‘Timeout’.

For species were gene duplication and loss were common, few single-copy orthogroups were identified with all species present (Figure 1 & Table 1). In particular, no single-copy orthogroups with all species present were identified in the Plants. This makes it impossible to infer a species tree using only single-copy orthologues. To overcome this problem, and allow us to infer a species tree using ASTRAL, NJst and concatenated multiple sequence alignments (CMSA), we relaxed the data selection criteria allowing selecting orthogroups that were single-copy for a proportion of species. This method allowed a smaller proportion of species to be present and single-copy in an orthogroup if it resulted in a proportionally greater increase in the number of orthogroups that could be used (Supplemental File S1 Figure S2, Supplemental File S1 Table S4). In all cases, only the single-copy orthologues in these orthogroups were used for species tree inference and multiple copy genes were removed from the alignment. This made it possible to infer a species tree with CMSA, ASTRAL and NJst for the plants dataset.

This relaxed data selection criterion improved the accuracy of CMSA and ASTRAL on average across the 12 datasets used in this study and thus these trees were used for CMSA and ASTRAL when comparing against STAG. STAG and Guenomu have no requirement for single-copy orthogroups and so this orthogroup selection method was not required for these methods.

### Software availability

STAG is written in python. The source code and precompiled binary are available in the Supplemental material (Supplemental File S2) and at https://github.com/davidemms/STAG. The software is operated via command line interface and can be used on Linux operating systems.

## Disclosure declaration

The authors declare no competing interests.

## Acknowledgements

SK is a Royal Society University Research Fellow. This work was supported by the European Union’s Horizon 2020 research and innovation programme under grant agreement number 637765. The authors would like to acknowledge the use of the University of Oxford Advanced Research Computing (ARC) facility in carrying out this work. http://dx.doi.org/10.5281/zenodo.22558

